# PDFGRα^+^ Stromal Cells Promote Salivary Gland Proacinar Differentiation Through FGF2-dependent BMP7 Signaling

**DOI:** 10.1101/2021.11.19.469144

**Authors:** Nicholas Moskwa, Ayma Mahmood, Deirdre A. Nelson, Amber L. Altrieth, Paolo Forni, Melinda Larsen

## Abstract

3.0

Stromal cells can direct epithelial differentiation during organ development; however, these pathways remain poorly defined. FGF signaling is essential for submandibular salivary gland development, and FGF2 can regulate proacinar cell differentiation in organoids through autocrine signaling in stromal cells. We performed scRNA Seq and identified stromal cell subsets expressing *Fgf2* and *Fgf10* that also express *Pdgfrα*. When combined with epithelial cells in organoids, MACS-sorted PDGFRα^+^ cells sufficiently promoted proacinar differentiation. Gene expression analysis revealed FGF2 activates the gene *Bmp7* in the stroma. BMP7 could replace stromal signaling and stimulate epithelial acinar differentiation but not branching. However, in the absence of FGF2, pathway analysis revealed that the stromal cells differentiated into myofibroblasts. Myofibroblast differentiation was induced when we treated organoids with TGFβ1, which also prevented proacinar differentiation. Conversely, FGF2 reversed TGFβ’s effects. Dissecting pathways driving acinar differentiation will facilitate development of regenerative therapies.

**Summary Statement:** Embryonic salivary glands contain multiple stromal cell populations. FGF2 maintains the stromal *Pdgfrα*^+^ population *in-vitro*. The PDGFRα^+^ stromal cells drive early epithelial secretory cell differentiation using BMP7.

## 4.0 Introduction

All developmental processes require precise cellular organization and coordination between many cell-types. The branching morphogenesis occurring in the salivary gland’s parenchymal epithelium requires instructions from embryonic mesenchymal cells (Sakakura et al., 1976; Wei et al., 2007). These stromal to epithelial interactions shape epithelial differentiation, instructing either secretory acinar or ductal cell fates. Functional salivary acini produce saliva and necessitates good oral health (Pedersen et al., 2018). However, acinar cells accumulate damage from diseases and treatments, including Type II diabetes, Sjögren’s syndrome, oral drug usage, as well as head and neck radiation (Jensen et al., 2014; B. Liu et al., 2012; Marmary et al., 2016; Plemons et al., 2014; von Bültzingslöwen et al., 2007; Weng et al., 2018; Yoo et al., 2014). Diseased and dysfunctional salivary gland treatments require understanding what molecular mechanisms underly these stromal-epithelial cell interactions driving secretory acinar cell differentiation.

Tissue recombination experiments first demonstrated the salivary gland’s mesenchymal cells impact the epithelium’s differentiation (Hoffman, 2002; Kusakabe et al., 1985; Sakakura et al., 1976; Wei et al., 2007). In the mouse submandibular salivary gland (SMG), fibroblast growth factor (FGF) signaling is a known essential mediator between the stroma and epithelium. FGF signaling drives many aspects of salivary gland development (Chatzeli et al., 2017; Hoffman, 2002; Lombaert et al., 2013; Makarenkova et al., 2009; Steinberg et al., 2005; Wei et al., 2007). The FGF growth factor family is a large and complex family with 23 known interacting members with at least the four FGF receptors and multiple isoforms (Yun et al., 2010). Salivary gland morphogenesis requires FGF signaling. Mice with FGFR2IIIb knocked-out exhibit abnormal salivary gland development (De Moerlooze et al., 2000; Hideyo et al., 2000). As FGFR2IIIb is expressed primarily in the epithelium, the epithelium’s failure at receiving a stromal signal stunts development. Similarly, when FGFR2IIIb receptor’s ligand, FGF10, is knocked-out the early gland’s development fails (Steinberg et al., 2005). This outcome reflects the reciprocal where the stromal cells cannot send the proper signal. The salivary gland’s connective tissue houses many cell populations, including the vasculature and nerves, that play similar roles (H. R. Kwon et al., 2017; Nedvetsky et al., 2014; Takebe et al., 2013, 2015).

FGF2, or basic FGF (bFGF), signaling is another growth factor implicated in acinar cell differentiation in the submandibular salivary gland (Hosseini et al., 2018). FGF2 is a paracrine signaling molecule that regulates many developmental processes including limb formation and wound healing (De Moerlooze et al., 2000; Martin, 1998). These processes require regulation across cell survival, migration, proliferation, and differentiation. In previous work, we developed salivary organoids made from E16 epithelial and stromal cells that produced FGF2-depenedent proacinar differentiation (Hosseini et al., 2018, 2019). These organoid experiments demonstrated that knocking down FGF2 specifically within the stroma inhibits the epithelium’s proacinar differentiation. Additionally, FGFR pharmacological inhibition in the organoids also prevented proacinar differentiation. However, FGF2 did not directly stimulate proacinar differentiation without the stroma. We demonstrated that FGF2 supports proacinar differentiation through the stromal cells. However, what epithelial differentiation signals the FGF2 was stimulating in the stroma was not addressed.

Single cell RNA sequencing (scRNA Seq) has revealed high levels of cellular heterogeneity within organs using hierarchical clustering (Enge et al., 2017; Hauser et al., 2020; O. J. Kwon et al., 2019; Lu et al., 2016; Macosko et al., 2015; McCarthy, Manieri, et al., 2020; Sekiguchi et al., 2020; Song et al., 2018; Zepp et al., 2017, 2021). These large datasets and their computational models have identified previously unknown cell types, confirming both known cellular communications driving development and exploring previously unknown pathways. In this study, we demonstrate that there are high levels of cellular heterogeneity within the salivary gland stroma during the proacinar cells’ differentiation. Moreover, we observed that FGF2 indirectly drives proacinar development through transcriptional changes regulating the stromal cell phenotype. *Pdfgrα* is one FGF2-controlled gene that is expressed in several stromal subpopulations. These PDGFRα^+^ stromal cells sufficiently promote epithelial proacinar differentiation. The FGF2 signaled stroma increases *Bmp7* expression, and BMP7 promotes proacinar differentiation. FGF2 also prevents the stroma’s differentiation into myofibroblasts and reverses TGFβ1-mediated proacinar de-differentiation in organoid cultures. These data demonstrate FGF2’s importance maintaining the stromal cell phenotype correlates with controlling proacinar cell differentiation.

## 5.0 Results

### 5.1 Identification of multiple salivary gland stromal cell subpopulations

As our previous studies showed SMG proacinar development requires stroma at the E16 developmental timepoint (Hosseini et al., 2018), we hypothesized that specific stromal subpopulations support proacinar differentiation. To address this hypothesis, we removed most epithelial cells, leaving behind enriched stromal cells from E16 SMGs. We then performed scRNA Seq on the enriched stromal cells. Stromal enrichment was necessary because prior studies showed that the salivary gland’s higher epithelial concentration lowers stromal cell representation (Hauser et al., 2020). The compiled scRNA data retained 9,165 total sequenced cells (Fig. S1).

The transcriptional subpopulations were modeled using Seurat’s principal component analysis (PCA) followed by k-nearest neighbor clustering and Louvain algorithms (Satija et al., 2015). We visualized the modeled subpopulations using graph-based dimensional Uniform Manifold Approximation and Projection (UMAP) reductions. Based on cells expressing epithelial cell adhesion molecule, *Epcam,* we sequenced 1,195 epithelial cells and 7,970 non-epithelial cells. Within the non-epithelial cells, we identified many cell-types residing within the connective tissue compartment (Fig. 1A, 1B & Fig. S1). *Col1a1* expression indicated we sequenced 3,790 stromal cells. The remaining 4,180 cells included endothelial cells, lymphatic endothelial cells, immune cells, neurons, Schwann cells, and erythrocytes, as defined by marker gene expression. We segregated and reanalyzed the epithelial populations looking for subpopulations. The remodeling recomputed the unsupervised clustering in Seurat (Satija et al., 2015) and was followed with supervised naming assignments based on multiple marker gene’s expression (Fig. 1C). This analysis yielded 9 epithelial subpopulations, including known epithelial sub-types such as, *Krt5*^+^ and *Krt19*^+^ ductal cells and *Epcam*^+^ and *Acta2^+^* myoepithelial cells (Hauser et al., 2020; Song et al., 2018). Populations expressing cell cycle genes were identified and labeled with “CC.” Additionally, we found that the two distinct proacinar populations reported existing at post-natal day 1 (P1), expressing either *Smgc*^+^ (MUC19) or *Bpifa2*^+^ (PSP) (Hauser et al., 2020), are detectable as early as E16. Our epithelial cell clustering was thus consistent with prior studies but detected epithelial differentiation events at the E16 time point.

**Figure 1:**
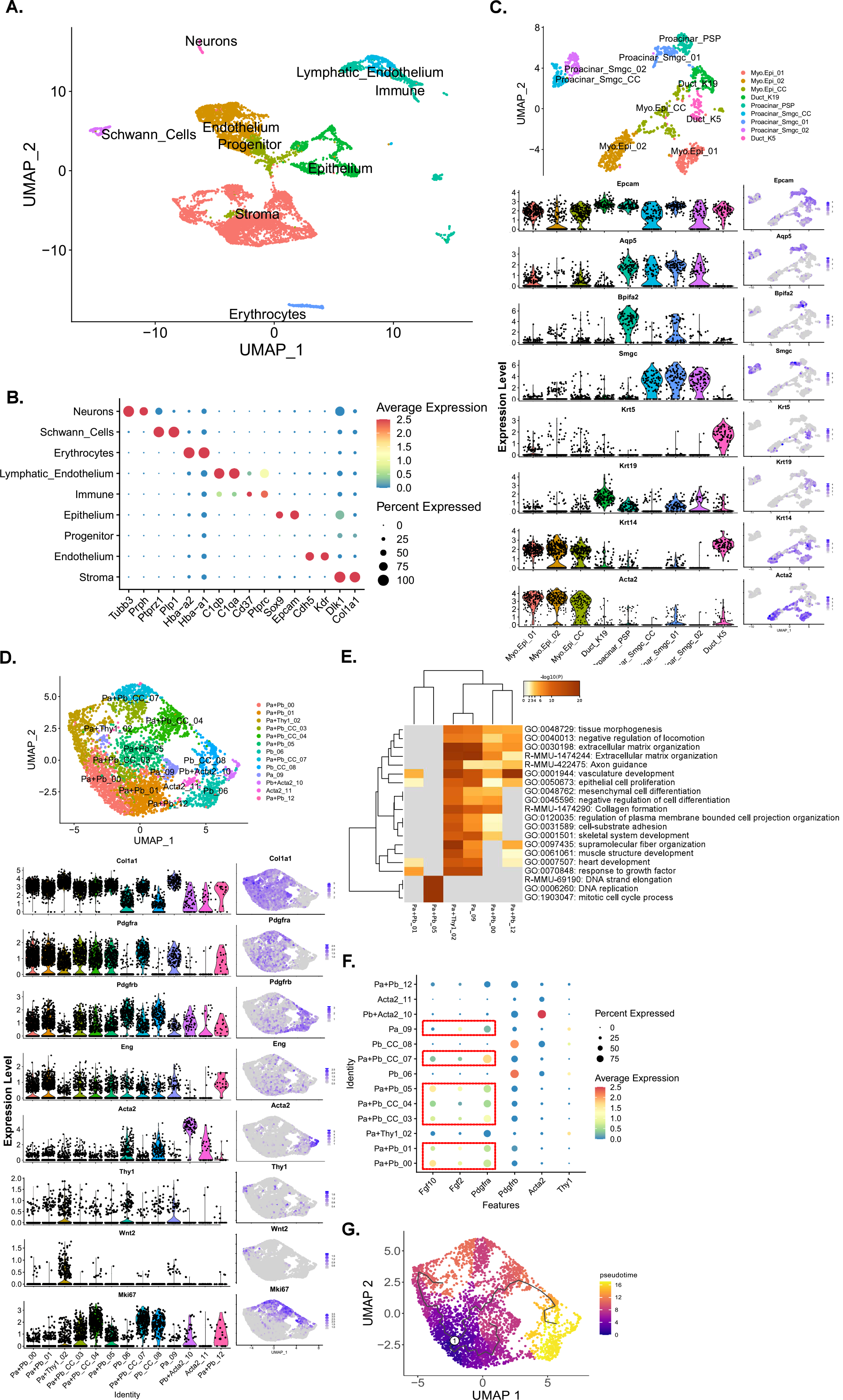
scRNA sequencing shows distinct stromal subpopulations in E16 SMG. A) Seurat generated UMAP plot using scRNA data from enriched stromal E16 SMG. Supervised clustering and gene expression instructed cell-type labeling. B) Dotplot showing 2 marker genes per cell-type and their corresponding expression levels. Gene expression informed cell cluster labeling. C) UMAP and violin plots generated subseted epithelial data. D) UMAP and violin plots generated subseted stromal data. E) Metascape analysis of non-proliferative *Pdgfrα*^+^ stromal cells. F) Dotplot showing only stromal gene expression. A correlation between *Pdgfrα* with *Fgf2* and *Fgf10* expression (Red boxes). G) Stromal subset data interpreted using pseudotime modeling. *Pdgfrα & Pdgfrβ* cells were chosen as the root node. This differentiation projection shows progression from *Pdgfrα & Pdgfrβ* cells through cell cycle to reach the *Acta2*^+^ population, but directly shifts into the Thy1^+^ population.

We segregated and re-clustered the stromal population to classify stromal cell subpopulations. This analysis modeled 13 previously undefined subpopulations from the 3,790 sequenced cells (Fig. 1D). Proliferative subpopulations accounted for 4 of the stromal clusters (SC_3, 4, 7, and 8), while the other 9 subpopulations are classifiable based on known stromal identity markers: *Pdgfrα*, *Pdgfrβ*, *Acta2*, *Thy1*, and *Wnt2* (Contreras et al., 2019; Dominici et al., 2006; Karpus et al., 2019; Kishi et al., 2018; Li et al., 2018; Santini et al., 2020; Zepp et al., 2017, 2021). Both Platelet Derived Growth Factor Receptor Alpha (*PDGFRα*/CD140a) and PDGFR Beta (*PDGFRβ*/CD140b) are especially interesting because they have been shown to play specific roles in organ epithelial differentiation, wound healing, and disease phenotypes (Kinchen et al., 2018; Kramann et al., 2015; Kuppe et al., 2021; Li et al., 2018; N. Liu et al., 2019; Ramachandran et al., 2019; Santini et al., 2020). PDGFRα is expressed in the neural crest-derived SMG mesenchyme and regulates early branching morphogenesis by stimulation of FGFs (Hoffman, 2002; Morikawa et al., 2009; Sakakura et al., 1976; Soriano, 1997; Steinberg et al., 2005; Yamamoto et al., 2008). We annotated the *Pdgfrα^+^* subpopulations’ transcriptomes using Metascape (Zhou et al., 2019) and found transcriptional landscapes matching the stroma’s traditionally understood functions. These *Pdgfrα^+^* clusters showed differing expression in extracellular matrix organization, vasculature development, and supramolecular fiber organization (Fig. 1E). We evaluated subpopulations expressing two growth factors with known importance in proacinar differentiation, *Fgf2* (Hosseini et al., 2018) and *Fgf10* (Chatzeli et al., 2021; Patel et al., 2007; Wells et al., 2013) (Fig. 1F). Of the 9 *Pdgfrα*^+^ subclusters, 7 express *Fgf2* and *Fgf10* at varying levels, suggesting that these specific stromal subpopulations participate in driving proacinar differentiation.

To explore the developmental relationship between heterogeneous stromal cell-types, we modeled cell differentiation trajectories in pseudotime (Fig. 1G). Using Monocle DDRTree (Qiu et al., 2017) we inferred trajectories from the E16 scRNA Seq transcriptomes. We set the root node as the non-cell cycling *Pdgfrα^+^* subpopulation because the total stromal population was reported arising from the *Pdgfrα^+^* migratory cranial neural crest (Morikawa et al., 2009). This trajectory analysis shows three branches. One branch that directly links the *Pdgfrα^+^* root subpopulation with the *Wnt2^+^*&*Thy1^+^* subpopulation, a second branch leads to another *Pdgfrα^+^* subpopulation that expresses *Pdgfrα*, and the third branch showing the *Pdgfrα^+^* subpopulation transitioning through the cell cycle before becoming the *Pdgfrβ^+^*&*Acta2^+^* subpopulation. This analysis predicts that the *Pdgfrα*^+^ progenitor population can differentiate along three trajectories becoming *Wnt2^+^*&*Thy1^+^*, *Pdgfrβ^+^*&*Acta2^+^*, or continuing as *Pdgfrα^+^* cells.

### 5.2 FGF2 promotes proacinar while inhibiting ductal differentiation

As organoids can be used to evaluate stromal-epithelial cell signaling, we examined the salivary gland organoid’s changes in epithelial gene expression when responding to FGF2. We produced organoids containing E16 SMG epithelial and stromal cells grown with or without FGF2, as performed previously (Hosseini et al., 2018, 2019). Our previous work demonstrating that FGF2 in organoids doesn’t directly promote epithelial to proacinar differentiation (Hosseini et al., 2018, 2019), made us question if the epithelium’s differentiation state change is detectable at the transcriptional level. We separated the organoid epithelial cells from the stromal cells and identified epithelial differentiation transcriptomes using Clariom S microarrays. We detected a clear separation between the control- and FGF2-treated epithelial cells in the PCA’s first principal component (Fig. S2C). We generated a 2-fold change heatmap, based on signal, and noted specific acinar and ductal differentiation markers showing contrasting signal magnitudes respectively between the FGF2 and control conditions (Fig. 2A). Gene set enrichment analysis (GSEA) identified global differences (Subramanian et al., 2005) showing acinar and ductal genes enriching in FGF2 and control conditions respectively, but not myoepithelial genes. All three gene-sets were derived from post-natal day 8 SMG epithelial scRNA Seq data (Song et al., 2018). FGF2 enriching the acinar phenotype was expected based on our prior work (Hosseini et al., 2019, 2018). However, organoids grown without FGF2 enriched for ductal genes and neither organoid had meaningful myoepithelial gene enrichment (Fig. 2B). Using immunocytochemistry (ICC), we confirmed that organoids lacking FGF2 increased the ductal marker, KRT7, and was inversely correlated with the acinar marker, AQP5 (Fig. S2A, B). Together, these results indicate FGF2 promotes a proacinar phenotype detectable at the transcript level. These data further indicates that in the absence of FGF2 the epithelial cells undergo a ductal differentiation.

**Figure 2:**
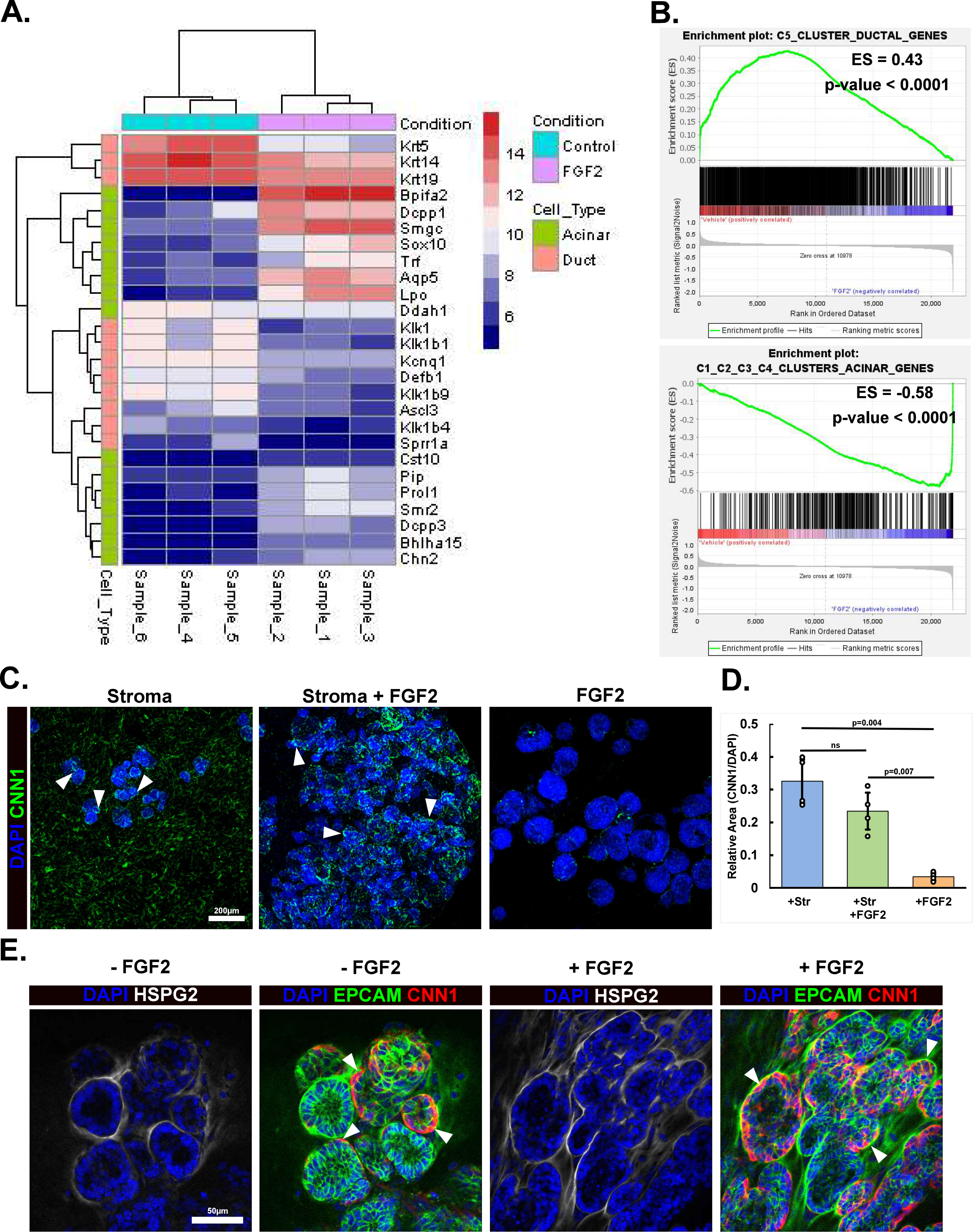
Proacinar, ductal and myoepithelial phenotypes are controlled in salivary gland organoids by FGF2 and stroma. A) Transcriptome heatmap showing enrichment of specific acinar (green) and ductal (salmon) genes with FGF2 (lavender) and control (cyan) organoid culture conditions, respectively. Log two-fold scale for gene expression is shown with red (higher) and blue (lower). B) GSEA plots depicting microarray differential gene expression lists compared with ductal and acinar gene lists. Plots show FGF2 cultured organoids enriched for acinar genes, while control organoids are enriched in ductal genes. Enrichment score (ES) and nominal p-value are provided. C) E16 SMG organoids grown with or without stroma and with or without exogenous FGF2 were immunostained to detect Calponin 1 (CNN1, green) with the nuclear stain DAPI (blue). Only organoids grown with stromal cells contain CNN1^+^ epithelium. Scale: 200 µm. D) Quantification of organoid area positive for CNN1 normalized to DAPI. Organoids made with stroma have significantly more myoepithelium; mean±s.d n=4 technical replicates, Single-factor ANOVA with post-huc Tukey test. E) Immunostained E16 SMG epithelial and stromal cell-containing organoids cultured with and without FGF2. An EPCAM^+^ (green) and CNN1^+^ (red) myoepithelial layer is present within in the epithelial compartment surrounded by HSPG2^+^ (white) basement membrane (arrowheads) both in the presence and absence of FGF2. Nuclei stained with DAPI (blue). Scale: 50 µm.

### 5.3 Stromal cells are required for myoepithelial cell differentiation independently of FGF2

To further specify FGF2’s contribution in epithelial differentiation, we searched for myoepithelium. As *in-vivo* SMGs contain proacinar and adjacent myoepithelium in bilayered acini, we looked for evidence that FGF2 regulates myoepithelial cell differentiation in the organoids. To assess a myoepithelial cell dependence on FGF2, organoids were grown for 7 days with or without stroma and with or without FGF2, then immunostained with the myoepithelial marker, Calponin-1 (CNN1). When stroma was present, CNN1 was detected in the organoids, but stromal deficient organoids had little to no detectable CNN1. Stromal containing organoids had similarly detected CNN1 levels independent of FGF2 presence (Fig. 2C, D). To confirm that CNN1 was expressed by epithelial cells, we grew epithelial-stromal organoids +/- FGF2 and performed ICC using CNN1 together with the epithelial marker, EPCAM, and the basement-membrane protein heparan sulfate proteoglycan 2 (HSPG2/perlecan). With high magnification confocal imaging, we confirmed that the EPCAM^+^ cells expressing CNN1 and were enclosed within the HSPG2^+^ basement membrane (Fig. 2E), as expected for myoepithelial cells. These data demonstrate that stromal cells are sufficient to achieve myoepithelial cell differentiation, independent of the FGF2’s presence.

### 5.4 FGF2 promotes stromal cell populations in organoids that mirror cell populations *in-vivo*

As we had previously determined that the stomal cells are the direct targets of FGF2 signaling in salivary gland organoids, we evaluated FGF2’s effects on stromal cell phenotype and compared the cell-types with IHC staining *in-vitro*. We first grew E16 stromal cells on glass coverslips or on porous polycarbonate filters in the presence or absence of FGF2. ICC showed that both the FGF2 and culture surface impacted marker expression. Stromal cells grown on glass with FGF2 showed a different cellular morphology than cells grown without FGF2. Although the PDGFRβ area was constant, there was a 3-fold increase in the area of cultured cells expressing PDGFRα in the presence of FGF2 (Fig. 3A, B). Additionally, THY1 expression was generally elevated in stromal cells grown with FGF2 growth conditions on glass (Fig. 3C, D). Independent of FGF2, the stroma’s organization was dependent on the culture surface. Although the stromal cells grew as a 2D monolayer on glass, when cultured on the filters, the stromal cells coalesced into 3D spherical aggregates, reminiscent of MSCs in a classical colony-forming assay (Friedenstein et al., 1970). Surprisingly, stromal cells grown on softer polycarbonate filters with FGF2 increased relative cell area positive for THY1 3-fold. In organoids treated +/- FGF2, the FGF2-dependent PDGFRα, and FGF-independent protein PDGFRβ expression in stroma was consistent with the 2D cultures (Fig. 3E, F). These data show stromal cell PDGFRα and THY1 protein levels are dependent on FGF2 signaling. To examine these cell populations *in-vivo*, we performed IHC on SMG tissue sections. We identified key stromal markers, PDGFRα, PDGFRβ, VIM, ENG, and THY1 (Fig. 3G) using IHC in E16 and adult SMGs. We detected stromal cell subpopulations expressing markers comparable with the E16 gland’s scRNA Seq data (Fig. 1C). This marker expression resembled FGF2 treated organoid stroma. The scRNA Seq showed a small THY1^+^ stromal subpopulation that was also Wnt2^+^ and PDGFRα^+^ (Fig 1D). We immunostained both endothelial and neuronal cells using PECAM1 and TUBB3, respectively, and identified THY1^+^ stromal cells residing in a neuro-vasculature niche in the adult SMG (Fig. 3G). Overall, we detected stromal cell subsets in E16 SMG corresponding with subpopulations modeled using scRNA Seq. Additionally, we found that with FGF2 treated organoids have stromal cell subpopulations more resembling E16 SMG stroma than stromal cells from untreated organoids. Interestingly, these stromal cell subpopulations remained present in adult glands, although at an overall lower proportion of the total cells. These data suggest that the *Pdgfrα^+^*subpopulations, including the *Thy1^+^* cluster, may support proacinar differentiation in organoids.

**Figure 3:**
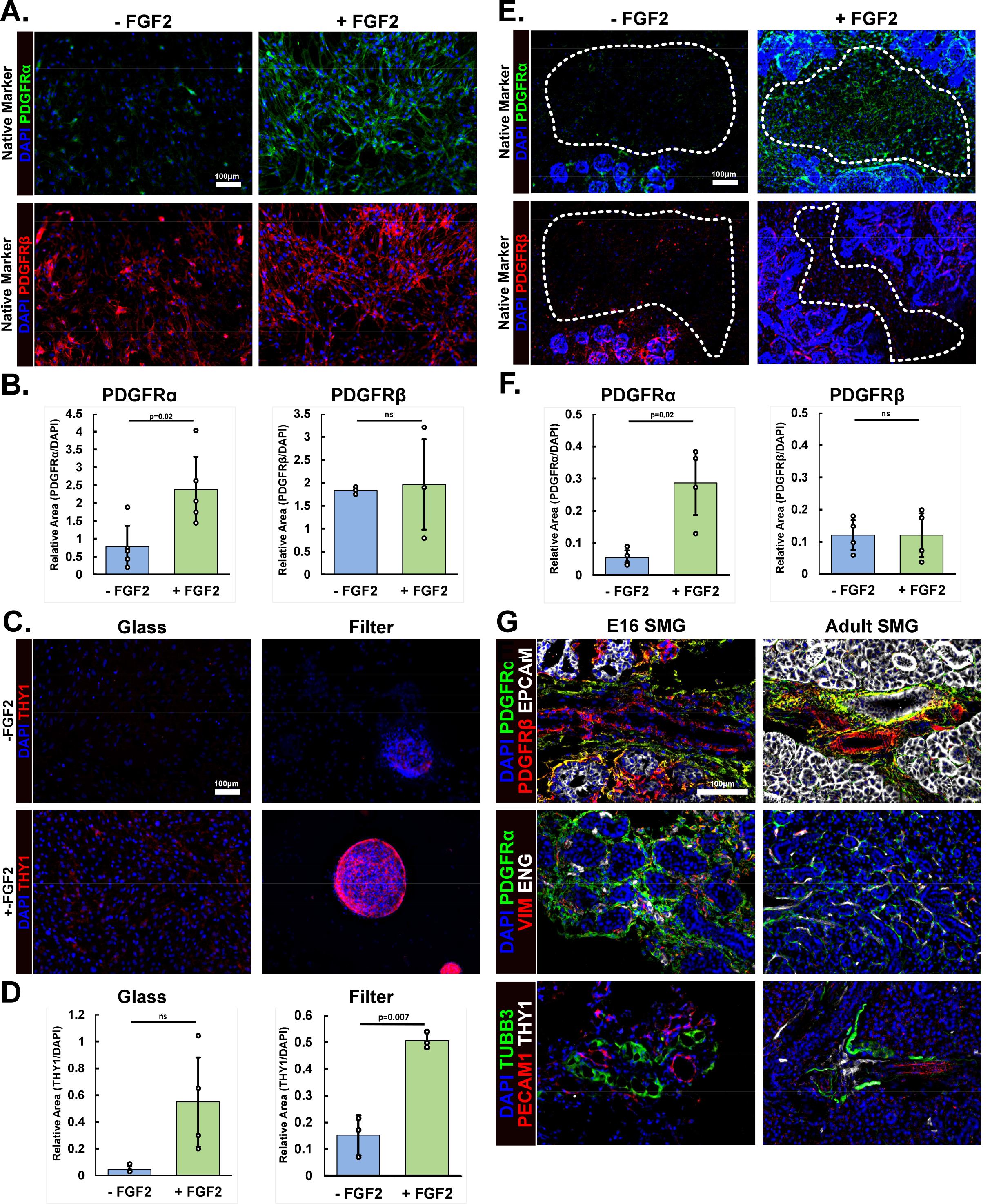
SMG Stromal cells cultured with exogenous FGF2 retain an *in-vivo*-like phenotype. A) E16 stroma cultured on coverslips in media containing or lacking FGF2. Cells positive for PDGFR (green) and PDGFR (red) increase when exogenous FGF2 is present, shown with DAPI (blue). Scale: 100 µm. B) Cell area quantification of PDGFRα and PDGFRβ normalized to DAPI. FGF2 increases stromal PDGFRβ-positive area 3-fold, but does not affect PDGFRβ-positive area significantly; mean±s.d n=5 and 3 experimental replicates, Two-tailed Unpaired Student t-test. C) E16 stroma cultured on coverslips or porous filters in media with or without FGF2 with ICC performed to detect THY1 (red) with DAPI (blue). Culture conditions and FGF2 signaling both affect E16 stromal THY1 expression. Only stromal cells cultured on filters with FGF2 organize into 3D clusters that express THY1. D) Area stain quantification of THY1 normalized to DAPI. A statistically relevant 3-fold increase in THY1 only occurs in cells cultured on the filter with FGF2 containing media; mean±s.d n=4 then 3 technical replicates, Two-tailed Unpaired Student t-test. E) Changes in stromal gene expression in the organoids. ICC was performed on E16 SMG epithelial and stromal organoids cultured with or without FGF2 containing media. PDGFRα (green) and PDGFRβ (red) staining show similar responses in organoids as with stromal only *in-vitro* culture. Only PDGFRβ is increased in organoid stroma (dotted white line) when FGF2 is present. PDGFRβ remains unchanged in either condition. F) Area stain quantification within organoids. PDGFRα/DAPI-positive area increases 5-fold in organoid stroma when FGF2 is added. β/DAPI-positive area does not change in organoid stroma; mean±s.d n=4 technical replicates, Two-tailed Unpaired Student t-test. G) Immunostaining for stromal markers performed on E16 and adult salivary gland cryosections. Both PDGFRα (green) and PDGFRβ (red) are expressed in E16 and adult SMG with singular and dual positive cells detected. Other stromal markers, Vimentin (VIM, red) and Endoglin (ENG, white), are also expressed in PDGFRα^+^ stromal cells. A rare stromal subset expressing THY1 (white) resides near neuronal TUBB3^+^ (green) and endothelial PECAM^+^ (red) cells within in a presumptive neural-vascular niche. Nuclei are labeled using DAPI (blue). Scale: 100 µm.

### 5.5 PDGFRα^+^ embryonic and adult stromal cells promote proacinar organoid differentiation with FGF2 dependency

We next questioned if the PDGFRα^+^ cells are sufficient to promote proacinar differentiation in organoids. We compared organoids generated using epithelial cells alone or epithelial cells grown together with either E16 SMG whole stroma or MACS-positively selected PDGFRα^+^ stromal cells. Epithelial only organoids treated with FGF2 demonstrated little to no PDGFRα^+^ stromal cell contamination (Fig. 4A, B). These FGF2-treated epithelial only organoids also showed very low levels of the proacinar maker, AQP5, and lacked budding structures (Fig. 4C, D). The FGF2-treated epithelial cell only organoids had a comparable morphology and differentiation state to the epithelial and whole stroma organoids grown without FGF2. The FGF2-treated organoids containing either PDGFRα^+^ stromal cells or total stroma had similar AQP5 expression and branched morphology (Fig 4C, D). Organoids made from PDGFRα^+^ stroma or total stroma showed a similar PDGFRα-positive area by day 7 (Fig 4A, B). Thus, we concluded that PDGFRα^+^ stromal cells, grown in the presence of FGF2, are sufficient to stimulate a budding proacinar organoid phenotype.

**Figure 4:**
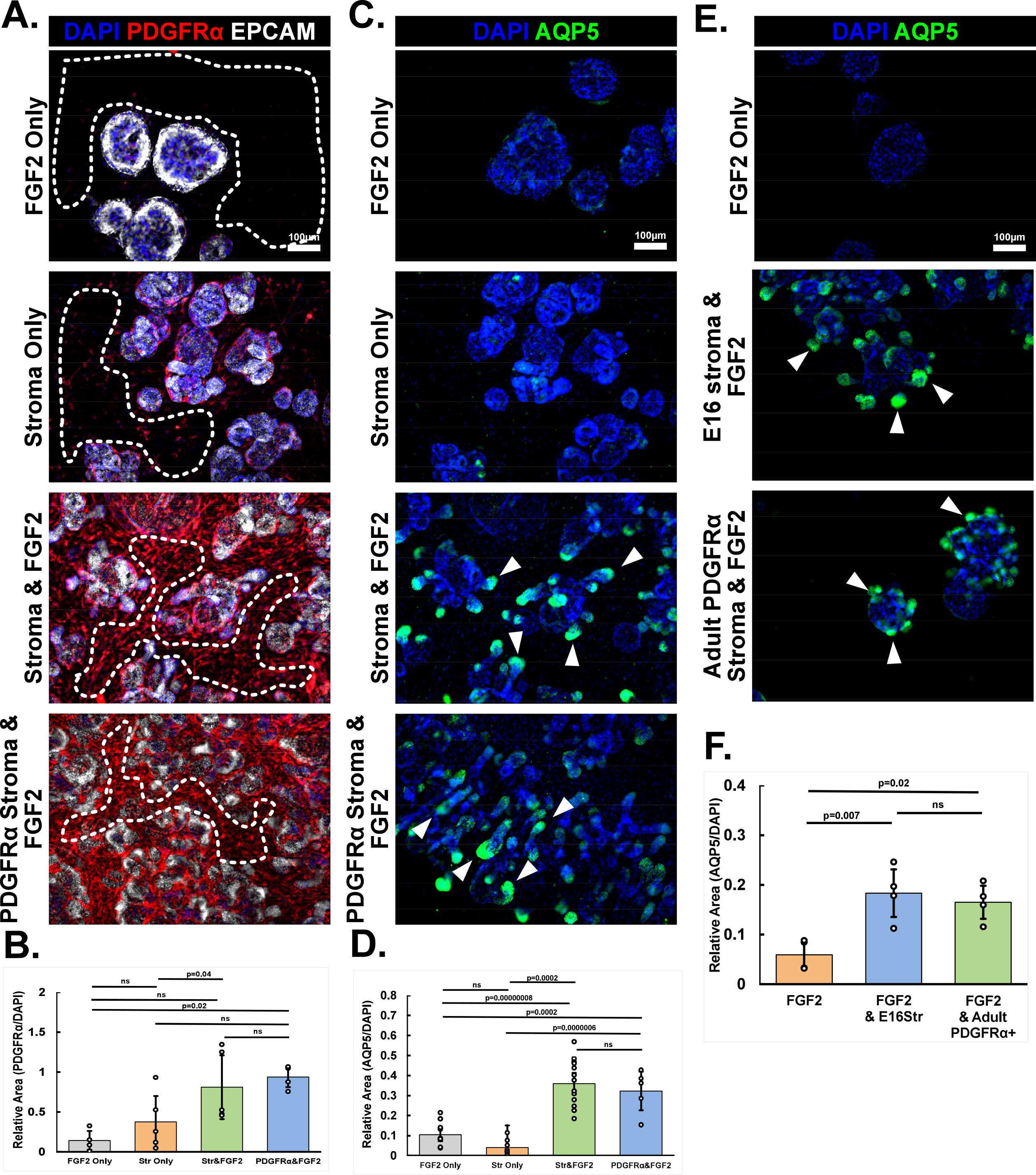
PDGFRα^+^ stromal cells promote proacinar organoids. A) E16 organoids created using epithelium cultured with: FGF2 and no stroma (FGF2 Only), stroma without FGF2 (Stroma Only), stroma and FGF2 (Stroma & FGF2) or with PDGFRα ^+^ MACS-isolated stroma and FGF2 (PDGFRα Stroma & FGF2). ICC for EPCAM (white) shows the epithelium, PDGFRα ^+^ (red) shows stromal subset, and DAPI (blue) shows nuclei. All condition form epithelial organoids, however stromal PDGFRα only increases when FGF2 is present. B) PDGFRα ^+^ positive area was quantified relative to DAPI. PDGFRα ^+^ MACS-selected stroma shows a >2-fold increase in PDGFRα similar to the total stroma when FGF2 is present; mean±s.d n=4, 5, 5, 4 experimental replicates, Single factor ANOVA with post-hoc Tukey Test. C) E16 organoids created using epithelium cultured with: FGF2 and no stroma (FGF2 Only), stroma without FGF2 (Stroma Only), stroma and FGF2 (Stroma & FGF2) or with PDGFRα ^+^ MACS-isolated stroma and FGF2 (PDGFRα ^+^ Stroma & FGF2). ICC for AQP5 (green) shows the proacinar epithelium and DAPI (blue) shows nuclei. AQP5 increases equally when either total stroma or PDGFRα ^+^ stroma are combined with FGF2. D) AQP5 positive area was quantified relative to DAPI. AQP5 increased when either total stroma or PDGFRα ^+^ stroma are in the presence of FGF2 containing media; mean±s.d n=11, 9, 14, and 5 experimental replicates, Single factor ANOVA with post-huc Tukey Test. E) Organoids created using E16 epithelium cultured in FGF2 media and combined with: no stroma, E16 total stroma, or adult PDGFRα ^+^ MACS-selected stroma. ICC showing proacinar epithelium using AQP5 (green) and DAPI (blue). The adult PDGFRα ^+^ stroma also support proacinar epithelium similarly to the total E16 stroma. F) APQ5 positive area quantified relative to DAPI. The organoids cultured with either E16 stroma or adult PDGFRα ^+^ stroma had a 3-fold higher AQP5^+^ relative area than organoids cultured without stroma; mean±s.d n=4 technical replicates, Single-factor ANOVA with post-hoc Tukey Test.

Since the stromal cell subpopulation appeared largely conserved in adult salivary glands, we tested the adult PDGFRα^+^ cell’s proacinar differentiation potency. We previously reported that total E16 stroma promotes adult proacinar differentiation *in-vitro* (Koslow et al., 2019) but did not examine adult stromal cells. We isolated PDGFRα^+^ stromal cells from the adult salivary gland using MACS and made organoids containing adult PDGFRα^+^ cells and enriched E16 epithelium. The adult PDGFRα^+^ stroma and E16 total stroma promoted similar AQP5 levels (Fig. 4E, F). We noted that the budded epithelial organoid regions were smaller and less distended during culture using adult rather than embryonic PDGFRα^+^ cells. These data demonstrate that adult PDGFRα^+^ stromal cells retain the proacinar promoting ability towards the E16 epithelium.

### 5.6 FGF2-stimulated stromal cells induce proacinar differentiation in organoids using BMPs

We wanted to identify FGF2-dependent changes in stromal gene expression driving proacinar differentiation. We extracted total RNA from stromal cells isolated from organoids grown for 7 days either with or without FGF2. We examined these RNA’s differential gene expression using Clariom D microarrays. The PCA’s first principal component showed over 50% of the variance and showed there were significant differences between the two conditions (Fig. 5A). We examined comprehensive transcriptomic changes that are FGF2-dependent using the Broads Institute gene-set database and GSEA (Subramanian et al., 2005). We observed that many stromal gene-sets enriched by FGF2 treatment were proliferation or E2F associated (Fig. 5B), which is expected given FGF2’s known participation in cell cycle (Krejci et al., 2004; Neary et al., 2005).

**Figure 5:**
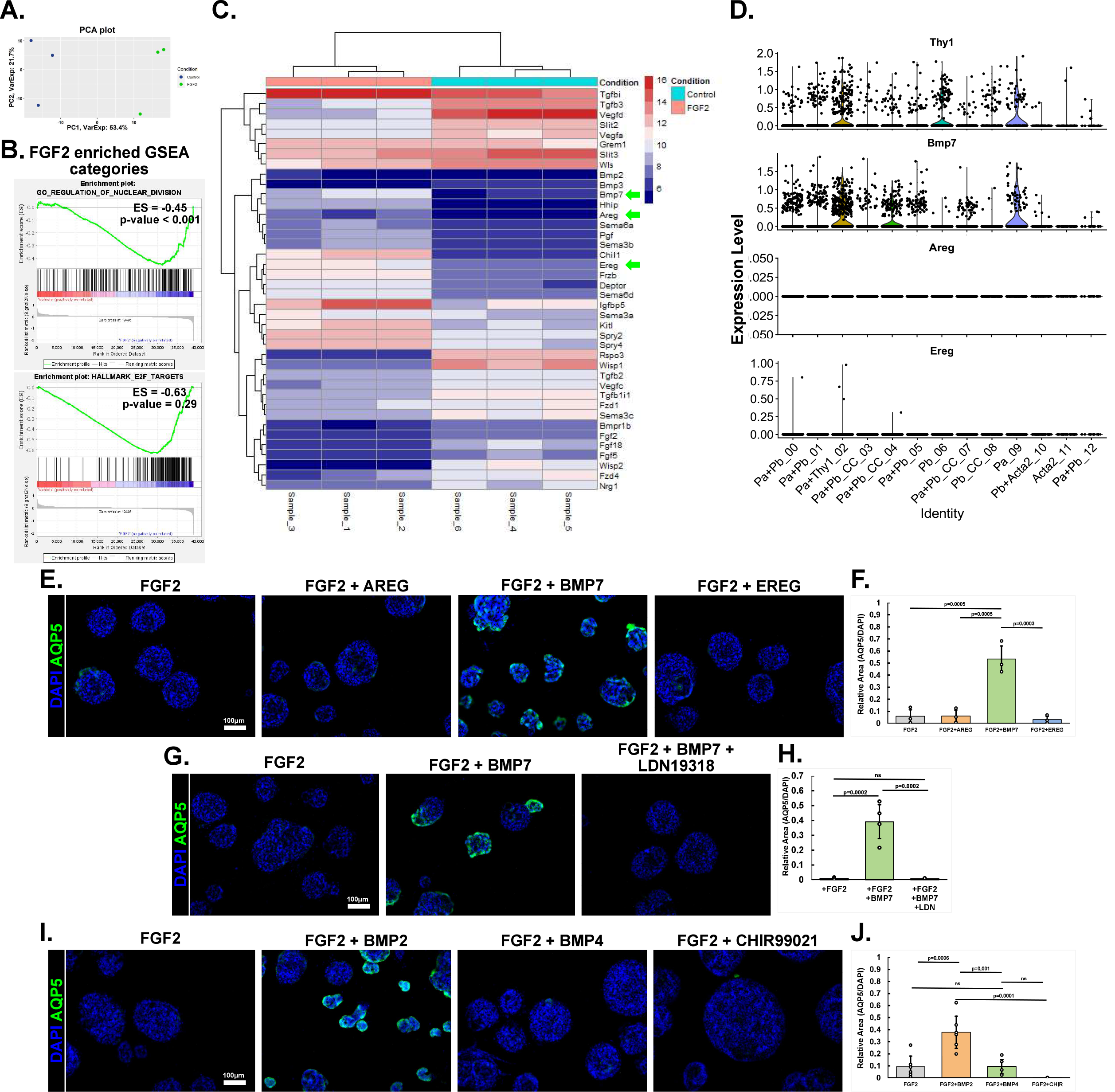
FGF2-dependent BMP signaling promotes epithelial proacinar differentiation. A) PCA plot comparing Clariom D microarray data generated from organoid stroma grown with or without FGF2. Each point represents one experimental replicate. More than half of the variance is described by the first principal component; n=3 experimental replicates. B) Enriched GSEA charts for microarray transcriptomes between enriched E16 organoid stroma with or without FGF2. Many of the stroma’s increased differenciated genes are associated with cell division. Enrichment score (ES) and nominal p-value are provided. C) Heatmap showing secreted factor changes from organoid stroma +/-FGF2. D) E16 scRNA Seq violin plots for stromal microarray genes. Only *Bmp7* transcription is present in E16 stromal subsets. E) ICC in epithelial only organoids cultured with FGF2 and either: AREG, BMP7, EREG. F) Epithelial organoid area positive for AQP5 normalized to DAPI. FGF2 and BMP7 cultured organoids show a significant 10-fold increase in relative stained area than FGF2 only cultured organoids; mean±s.d n=3 experimental replicates, Single-factor ANOVA with post-huc Tukey test. G) ICC in epithelial only organoids with FGF2 alone or in conjunction with either BMP7 or BMP7+LDN19318 (BMP inhibitor). H) Epithelial organoid area positive for AQP5 normalized to DAPI. LDN19318 significantly reduce organoids AQP5 relative area by 40-fold; mean±s.d n=4 experimental replicates, Single-factor ANOVA with post-huc Tukey test. I) ICC in organoids created using only E16 epithelium grown with media containing FGF2 and/or growth factors. Only BMP2 in combination with FGF2 promoted AQP5 expression. J) Epithelial organoid area positive for AQP5 normalized to DAPI. The FGF2 and BMP2 cultured organoids showed a significant 4-fold increase in relative stained area compared with FGF2 only cultured organoids; mean±s.d n=6, 6, 5, 4 experimental replicates, Single factor ANOVA with post-hoc Tukey Test.

To identify FGF2-regulated stromal genes that may promote the epithelium’s proacinar cell differentiation, we targeted the microarray’s growth factors that were differentially expressed.

From this list we selected three genes having a greater than 2-fold difference in expression +/- FGF2: *Areg*, *Ereg* and *Bmp7* (Fig 5C). *Areg* and *Ereg* are EGF homologs and EGF is a known growth factor promoting salivary gland development (Mizukoshi et al., 2016). Prior research illustrated BMP’s mixed contributions towards promoting salivary gland branching morphogenesis (Hoffman, 2002; Steinberg et al., 2005), but epithelial cell differentiation was not addressed. However, in intestinal organoids, hair follicles, and lungs, BMPs induce epithelial cell differentiation (Lu et al., 2016; McCarthy, Kraiczy, et al., 2020; Zepp et al., 2017). In the scRNA Seq modeled stromal subpopulations (Fig. 5D), both Areg and Ereg had low to no expression detected. However, *Bmp7* was expressed lowly in many *Pdgfrα*^+^ clusters and was surprisingly high in the *Thy1*^+^ cluster. We tested these three factors using organoids lacking stroma that were cultured in FGF2-containing media and found that only BMP7 induced AQP5 (Fig. 5E, F). However, we also noted that the organoids’ budding was low. We validated that BMP signaling in the epithelium is necessary for proacinar marker expression using a BMP inhibitor. Epithelial only organoids were treated with FGF2 and BMP7-containing media including a selective BMP receptor inhibitor, LDN19318, which targets primarily ALK2 and ALK3. Epithelial organoids grown with FGF2, BMP7, and LDN19318 had similar characteristics as the FGF2 only cultured organoids and no proacinar marker expression (Fig. 5G, H). As BMP2 was another FGF2 induced stromal protein and BMP4 is a signaling homolog to BMP2 and BMP7 (Bandyopadhyay et al., 2006; Choi et al., 2012; Kim et al., 2019), we also exposed FGF2-treated epithelial organoids to BMP2 or BMP4. BMP4 did not induce AQP5 expression. However, BMP2 did induce proacinar AQP5 expression (Fig. 5I, J) with similar levels of AQP5 as BMP7-treated organoids. We also used the WNT activator, CHIR99021, to simulate a cell cycle promotor analogous with the EGF homologs. As expected, CHIR99021 did not induce AQP5 expression but did create larger organoids. These data suggest that BMP7 and BMP2 are FGF2-dependent, stromal-produced, paracrine-acting factors promoting proacinar differentiation in SMG organoids.

### 5.7 FGF2 inhibits and reverses myofibroblast differentiation in salivary gland organoids

Based on the increased CNN1 expression in the stroma and reduced PDGFRα staining in organoids in the absence of FGF2, we questioned if lacking FGF2 phenotypically changes stromal cells. We examined the Clariom D microarray’s gene-sets enriched in the stroma grown without FGF2 (Fig. 6A). Three of the highest significantly enriched gene-sets, identified by GSEA, were for myogenesis, extracellular matrix, and TGFβ signaling. All three gene-sets align with myofibroblast characteristics, we hypothesized that without FGF2, the stromal cells differentiate into myofibroblasts. Myofibroblasts are known injury and fibrotic disease contributors that respond by increasing extracellular matrix deposition (Contreras et al., 2019; Kramann et al., 2015; Kuppe et al., 2021; Li et al., 2018; Nagaraju et al., 2019). We generated a heatmap for core myofibroblast genes that showed all genes were expressed relatively higher in control organoid stroma (Fig. 6B). Based on the scRNA Seq trajectory analysis (Fig 1F), the E16 stromal cells have the differentiation potential to express the classical myofibroblast gene smooth muscle alpha actin (αSMA/*Acta2*). Furthermore, we used GSEA to determine which scRNA Seq subpopulation best matched the control stroma’s transcriptome (Fig. 6D). The GSEA showed that the *Pdgfrβ*^+^ and *Acta2*^+^ stromal cluster 10 (SC_10) genes best enriched the control organoid stroma’s transcriptome.

**Figure 6:**
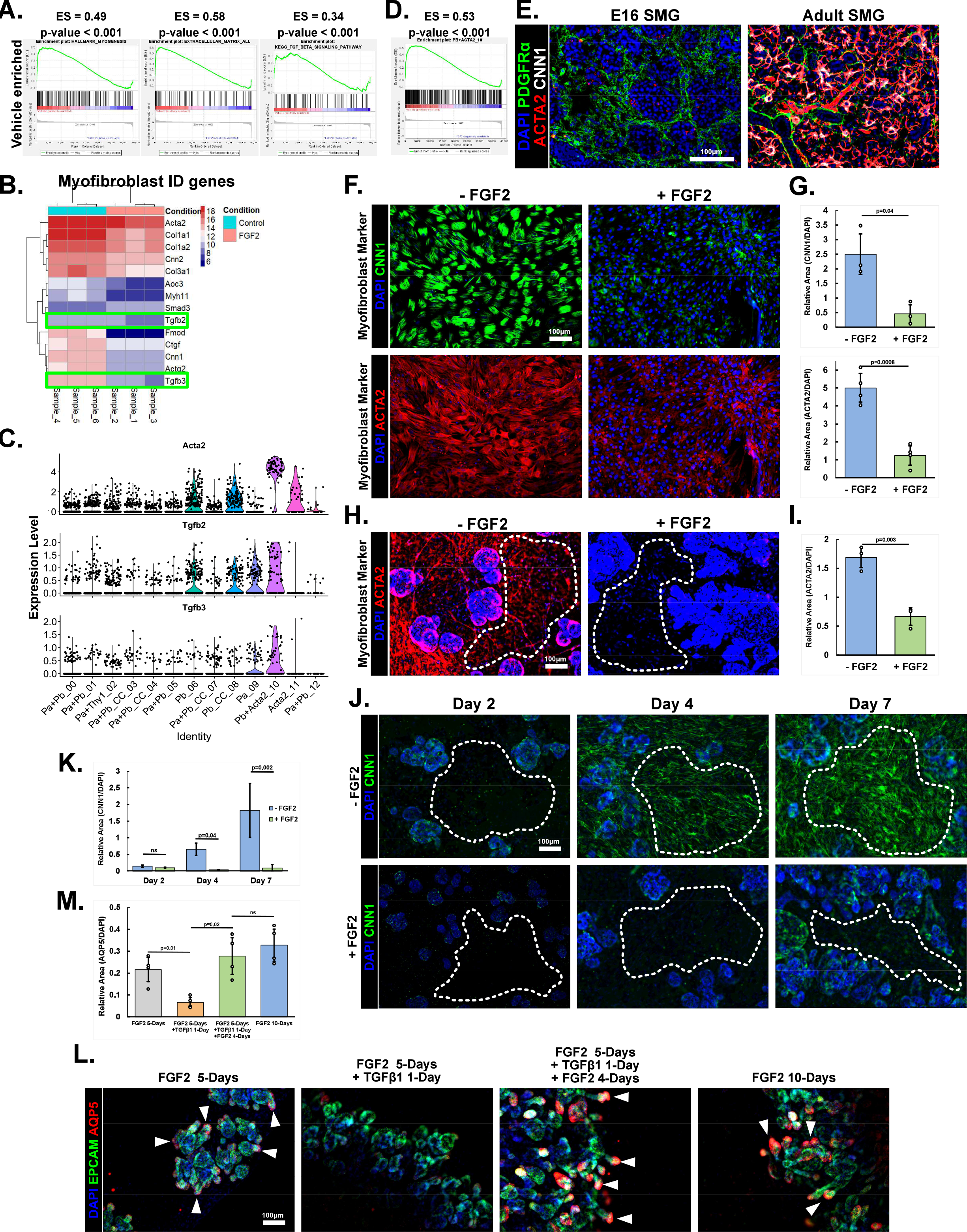
E16 Stromal cells cultured with exogenous FGF2 resist an inhibitory myofibroblast differentiation. A) E16 organoid stroma transcriptomes show myofibroblast enriched gene sets in control conditions. Enrichment score (ES) and nominal p-value are provided. B) Heatmap showing classical myofibroblast genes, including Smooth muscle alpha actin (*Acta2*), Calponin-1 (*Cnn1*), Transforming Growth Factor Beta-2 *Tfgβ*2) and Transforming Growth Factor Beta-3 *Tfgβ3*), are higher in E16 organoid stroma under control over FGF2 conditions. C) *Acta2, Tgfb2* and *Tgfb3* violin plots from E16 stromal scRNA Seq data. D) GSEA plot showing E16 organoid stroma’s control gene expression is similar to *in-vivo* smooth muscle cell phenotype described in scRNA Seq data. Enrichment score (ES) and nominal p-value are provided. E) ICC on E16 and adult murine salivary gland cryosections. PDGFRβ ^+^ stroma are not ACTA2^+^ or CNN1^+^ in the connective tissue cells from E16 or adult. The adult ACTA2^+^ or CNN1^+^ cells are likely myoepithelium. F) E16 stroma cultured on coverslips in the presence or absence of FGF2. Immunostaining shows both myofibroblast markers ACTA2 and CNN1 increase when the cells are cultured without FGF2. G) Quantification of myofibroblast markers area to DAPI area. A 5-fold increase occurs for myofibroblast marker ACTA2 and CNN1 in no-FGF2 cultured stroma; mean±s.d n=3 then 4 experimental replicates, Two-tailed Unpaired Student t-test. H) E16 epithelial and stromal organoids cultured with or without FGF2. The myofibroblast marker ACTA2 is much higher in stroma without FGF2. I) Quantification of ACTA2 area to DAPI area. ACTA2 is 2-fold higher in organoid stroma without FGF2; mean±s.d n=3 technical replicates, Two-tailed Unpaired Student t-test. J) E16 organoid developmental time-course on day 2, 4, and 7, using ICC for myofibroblast marker CNN1. The stroma gains the myofibroblast marker over time. K) Quantification of CNN1 area to DAPI area. The amount of CNN1 in stroma steadily increases as organoids grown without FGF2; mean±s.d n=3 technical replicates, Two-tailed Unpaired Student t-test. L) E16 epithelium and stromal organoids grown over 5, 6 and 10 days. FGF2 media applied first for 5 days to prevent myofibroblast phenotype and induce proacinar phenotype. Additional 1-day culture with TGFβ1 simulates a Myofibroblast response. The TFGβ decreases initial branching morphogenesis and AQP5 expression in epithelium. This short fibrotic induced environment is reversed when FGF2 is re-introduced for 4-days. M) Area stain quantification. The 1-day TFGβ1 introduction significantly reduces AQP5 expression by 3-fold. FGF2 re-introduction and culture time significant rebounded epithelial proacinar phenotype to match un-manipulated 10-day FGF2 organoids; mean±s.d n=4 technical replicates, Single factor ANOVA with a Tukey post-Hoc test.

**Figure 7:**
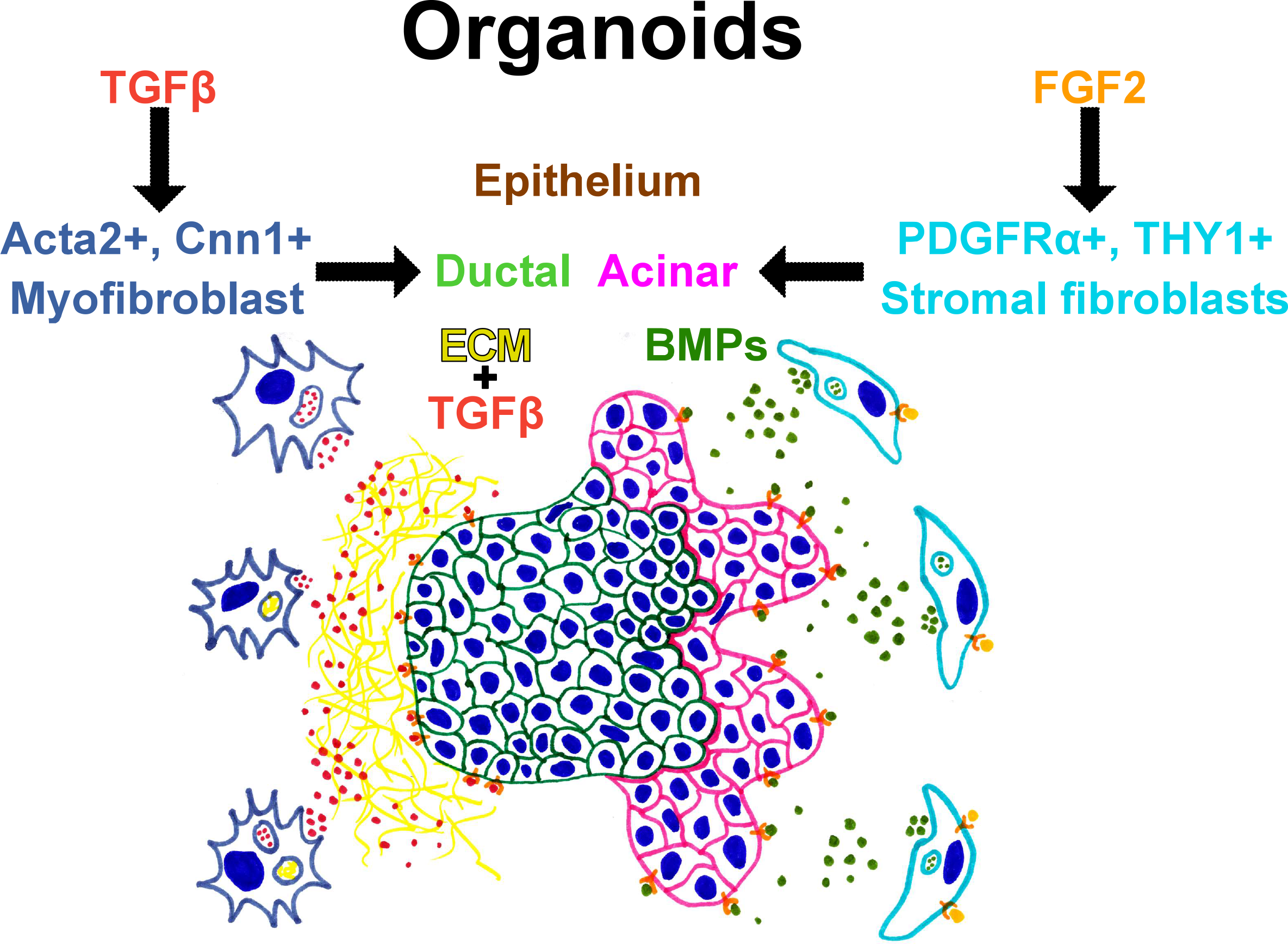
Schematic. A) TGFβ signaling signals stromal transdifferenciation into myofibroblasts. Myofibroblast appearence and TGFβ signaling correlates  with proacinar dedifferenciation. FGF2 signaling causes *in-vivo* stromal marker retention and BMP expression. These PDGFRα^+^ stroma specifically produce BMPs which cause proacinar differentiation. scRNA Seq data and organoid information inform how the cell are interacting in the salivary gland.

To determine if SMG stromal cells differentiate into myofibroblasts in the absence of FGF2, we cultured E16 stromal cells alone on coverslips with or without FGF2. The stromal cells grown without FGF2 had nearly a 5-fold elevation in the myofibroblast markers, ACTA2 and CNN1, relative to cells grown with FGF2 (Fig. 6F, G). To determine if the myofibroblast conversion is also occurring in the organoids, we examined ACTA2 and CNN1 expression in organoids grown +/- FGF2. We detected a similar myofibroblast transition in the organoids, but with a lower 3-fold change (Fig. 6H, I). A time-course was performed with time-points at 2, 4, and 7 days to explicitly examine a myofibroblast conversion in the organoids in the absence of FGF2. At day 2, there was no significant difference in CNN1 staining in the organoid stroma relative to FGF2 treatment (Fig. 6J, K). By day 4 there was noticeably more myofibroblast marker expression in the control organoids’ stroma, which continued into day 7. These data support the hypothesis that SMG stromal cells are transdifferentiating into myofibroblasts in the absence of FGF2.

Two mRNAs that were increased in no FGF2 organoid stroma were *Tgfβ2* and *Tgfβ3* (Fig. 6B), which are TGFβ related profibrotic cytokines that can induce myofibroblast differentiation and fibrosis (Avery et al., 2018; Nagaraju et al., 2019; Ó Hainmhire et al., 2019; Woods et al., 2015). Notably, the *Acta2^+^*&*Pdgfrβ*^+^ stromal cluster (SC_10) had appreciable *Tgfβ2* and *Tgfβ3* expression in the scRNA Seq-defined stromal cell subsets (Fig. 6C). Organoid experiments where TGFβ and FGF2 were switched showed a correlation between proacinar inhibition and myofibroblast induction (Fig. S3). We further tested whether TGFβ signaling inhibits proacinar differentiation by inducing organoid AQP5 expression after culturing five days with FGF2. Then FGF2 was washed out and replaced with TGFβ1-containing media. One day after TGFβ1 addition, the proacinar marker, AQP5, area decreased 3-fold (Fig 6L, M). Notably, this TGFβ1-induced AQP5 expression reduction was reversable when TGFβ1 media was washed away and FGF2-containing media reapplied. AQP5 re-expression occurred 4 days after FGF2 re-introduction. These FGF2-TGFβ1-FGF2-cultured organoids had the same relative AQP5 staining levels as organoids grown with FGF2 for 10 consecutive days. Together, these data indicate that TGFβ1 treatment inhibits proacinar organoid differentiation in a reversible fashion, and that FGF2 can re-induce proacinar differentiation following TGFβ1-induced proacinar inhibition caused by myofibroblasts.

## 6.0 Discussion

Even though it has been known for decades that stromal fibroblasts direct organ development, recent advances in scRNA Seq have illuminated the organ development contributions heterogenous stromal populations make. Here we performed scRNA Seq in the developing E16 SMG after stromal cell enrichment. In examining specific markers that describe functional stromal subsets in other organs, specifically, *Pdgfrα*, *Pdgfrβ*, *Acta2*, *Eng*, *Thy1* and *Wnt2* (Karpus et al., 2019; Kramann et al., 2015; O. J. Kwon et al., 2019; Lee et al., 2017; Li et al., 2018; Santini et al., 2020; Stzepourginski et al., 2017; Yang et al., 2017; Zepp et al., 2021; Zhao et al., 2017), we parsed subpopulations. UMAP regression and superimposed pseudotime models revealed that these stromal cells progress through a broad and continuous transcriptional landscape. Our pseudotime analysis interpretation speculates that these burgeoning subpopulations arise from a *Pdgfrα^+^* population that becomes multiple subpopulations. Other developmental scRNA Seq data employing pseudotime analysis suggests that embryonically defined epithelial subpopulations in the salivary gland and lung become more transcriptomically isolated over time (Hauser et al., 2020; Zepp et al., 2021). This transcriptomic isolation measures a cell’s differentiation progression, and it is not firmly established whether the stromal cells differentiate into distinct subpopulations as epithelial populations do or if their states remain more fluid. Recent studies reveal that in other organs, specific stromal subpopulations help create micro-niches that localize distinct epithelial identities into specific regions and mediate disease responses (Contreras et al., 2019; Li et al., 2018; McCarthy, Manieri, et al., 2020; Santini et al., 2020; Yang et al., 2017). Similarly, we identify cells in stomal cell subpopulations expressing *Pdgfrα* as cells that support proacinar cell differentiation in embryonic organoids.

Our developmental model suggests that the E16 stroma has multiple transcriptomic possibilities. The model predicts that these PDGFRα^+^ cells transition into smaller subpopulations upregulating either *Thy1* or *Acta2*. Our *in-vitro* studies, where stromal cells are cultured either with or without FGF2, validate that the PDGFRα^+^ cells can differentiate into at least two different subpopulations. These *in-vitro* culture studies revealed cells increasing levels of markers expressed by subpopulations modeled at the pseudotime’s end nodes. These data reflect that the stroma’s cell state, analogous to the epithelium, is dynamic at E16. The stroma and epithelia together complicate but direct each other’s signaling environment. Each cell-type’s transcriptomic and translational range highlights the organoid model’s usability in teasing apart complex cellular interactions. Our organoids inform this dynamic response between the epithelium and stroma. We see this cellular interaction happening where FGF2 maintains a stromal PDGFRα^+^ state mirroring *in-vivo* expression patterns and correlating with epithelial proacinar differentiation. These data suggest the stroma has a niche function for differentiating epithelium under these conditions. Studies have shown PDGFRα^+^ stromal cells have a transcriptomic plasticity that is important for regulating both developmental and injury responses (Contreras et al., 2019; Li et al., 2018). Here we provide data filling the gap between these two states where we explicitly define an inverse correlation between the stromal cells expressing myofibroblast markers when FGF2 is absent and the epithelium’s FGF2-induced proacinar differentiation state.

Using organoids, we determined which subpopulations direct proacinar differentiation. The neural crest markers’ (Morikawa et al., 2009) co-expression with proacinar signaling growth factors (Chatzeli et al., 2021; Hosseini et al., 2018; Patel et al., 2007; Wells et al., 2013) suggests that PDGFRα^+^ cells are potentially key contributors. Others have highlighted a theme where PDGFRα stromal expression aligns with functional dexterity (Contreras et al., 2019; Li et al., 2018; Santini et al., 2020). Our organoid experiments showed that these PDGFRα^+^ cells can recapitulate the total stroma’s function in promoting proacinar differentiation. Organoids containing embryonic cells and adult PDGFRα^+^ stroma demonstrated a functional maintenance in the stromal cell populations. The stroma’s continued instructiveness but proportional decline during development may reflect a switch in the epithelial dependency away from stromal paracrine signaling into homeostatic autocrine signaling. As the epithelial cells reach functional independency and maturity, the stroma may no longer be required at high density unless an insult occurs. The stromal to epithelial cell proportions vary from organ to organ and may provide a quantitative readout about the *in-vivo* recovery potential a given organ has.

Our study additionally suggests that the stromal state effects the epithelial differentiation decision into ducts or acini. FGF2 stimulated stromal PDGFRα and *Bmp* expression. The BMP signaling can direct the epithelium towards a proacinar rather than a ductal state. The organoid experiments in which only epithelial cells and growth factors are included suggests a separation in proacinar differentiation and branching morphogenesis. This idea is in line with older studies showing BMP7 does not contribute to branching morphogenesis (Hoffman, 2002; Steinberg et al., 2005), and a new study further advancing that cell to matrix adhesions direct branching morphogenesis (Wang et al., 2021). Stromal BMP-directed epithelial differentiation may also be a theme. A similar PDGFRα^+^ cell induced BMP gradient was recently shown to control differentiation progression in the small intestinal crypt (McCarthy, Kraiczy, et al., 2020), and the small intestinal PDGFRα^hi^ telocytes and our PDGFRα^+^ salivary stroma may be functionally and transcriptomically similar. This BMP shift contrasts second effect FGF2 is having in the salivary organoids. The FGF2 signaling is also preventing the PDGFRα^+^ cells from differentiating into ACTA2^+^ myofibroblasts. This myofibroblast phenotype produces at least one known branching morphogenesis inhibiting growth factor, TGFβ (Hall et al., 2010; Woods et al., 2015). TGFβ can inhibit proacinar differentiation and is likely promoting ductal differentiation, as previously reported (Iwano et al., 2002; J. Liu et al., 2016; Rastaldi et al., 2002; Rees et al., 2006; Termén et al., 2013). This BMP and TGFβ signaling axis may be one link between developmental maturation and injury recovery. Where TGFβ is important for early salivary gland development (Jaskoll & Melnick, 1999) and is co-opted during injury for re-purposing cells. Then BMPs used to differentiate cells during development (Bandyopadhyay et al., 2006; Lu et al., 2016; McCarthy, Kraiczy, et al., 2020; McCarthy, Manieri, et al., 2020) may also be used to re-differentiate adult cells post-injury.

Evaluating stromal heterogeneity remains a new and difficult frontier. Organs with lower stromal cell proportions, such as salivary glands (Hauser et al., 2020), require enrichment steps capturing enough cells to perform informative RNA Sequencing and hierarchical clustering. Our research suggests that the adult PDGFRα^+^ stroma has potential therapeutic relevance. However, more studies evaluating the evolution of stromal cell subpopulations at later developmental stages and in disease models will be needed to understand if the stroma’s developmental signaling is involved in disease progression. Our research here does demonstrate the advantage of evaluating predictions made with scRNA Seq using organoid models. Organoids offer the plasticity to test the many ideas scRNA Seq data generates at the cellular and signaling axes. Overall, understanding the stromal complexity in all organs, including the salivary gland, will create better *in-vitro* models that inform cell-therapy avenues.

## 7.0 Materials and Methods

### 7.1 Mouse submandibular gland cell isolation and enrichment

Embryonic day 16 (E16) timed-pregnant CD-1 female *Mus musculus* were ordered from Charles River Laboratories. First the E16 embryos, and then the submandibular glands (SMGs) were dissected out following animal protocols approved by the University at Albany Institutional Animal Care and Use Committee (IACUC). SMG removal involved slicing the mandible with sharp scalpels, then removing the glands using sterile forceps under a dissecting microscope. SMGs were micro-dissected in with 2x collagenase/hyaluronidase (StemCell Technologies #7912) diluted using 1xPhosphate Buffered Saline (PBS) (Life Technologies #70011-044). The solution was further diluted, and SMGs further dissected into lobules, when an additional volume of 1.6 U/ml of dispase II (Life Technologies, #17105041) was added creating a final concentration of 0.8 U/ml dispase II. The lobules were further broken-down after a 30-minute incubation at 37°C followed by trituration, generating epithelial clusters and single stromal cells. Enzymatic activity was quenched with 10% fetal bovine serum (FBS) (Life Technologies #10082147) in 1:1 DMEM/F12 (Life Technologies #11039047) supplemented with 100 U/mL penicillin and 100 mg/mL streptomycin (PenStrep) (Life Technologies #15140122). Cell populations were separated by gravity sedimentation for approximately 10 minutes until the large clusters formed a loose pellet. Epithelial clusters were enriched with two additional gravity sedimentations. Any DNA-induced clumping was stopped using 0.05 mg/mL DNase I (StemCell Technologies #07900) during the second gravity sedimentation. The stromal cell population was enriched by filtration through 70 µm (Falcon #087712) and 40 µm cell strainers (Fischer Scientific #22363547). Filtered stroma were pelleted at 300xG for 8 minutes and the buffer was replaced using DMEM/F12 containing 10% FBS and Pen-Strep.

Organoids requiring further epithelial cell enrichment, enrichment was performed as previously described (Hosseini et al., 2018, 2019). Epithelial enrichment involved dislodging any attached stroma by plating the clusters into a 35 mm tissue culture dish and incubating at 37°C for 2 hours. Stromal cells were removed from epithelial clusters using two washes with centrifugation at 10xG for 1-minute, supernatant removal, and resuspension in medium.

### 7.2 E16 Stroma Single Cell RNA sequencing

E16 SMG stromal cells were prepared as described above using both glands from 14 E16 pups excised from 2 pregnant moms. The live to dead cell ratios and cell numbers were evaluated with Trypan-blue (Gibco #15250-061) staining and a hemocytometer. The preparation had >80% viable cells and were frozen down in cryovials using 90% FBS + 10% Dimethyl Sulfoxide (Amresco #200-664-3). Cells were frozen at -1°C/minute in a 100% isopropanol freezing chamber overnight at -80°C. Frozen cells were stored in liquid-nitrogen chamber prior to shipping overnight on dry ice. Cellular recovery goal for library preparation was 10000 cells. Single cell RNA library preps were created and sequenced by Singulomics (Brooklyn, NY).

### 7.3 Computational Analysis of scRNA sequencing data

Bcl2fastq and CellRanger v4.0 assembled and counted sequence data. Data files were imported using SEURAT v3.1.5 in R v3.6.3 (R Core Team, 2021) & R Studio v1.2.5042. Data clusters were calculated following the default pipeline (Satija et al., 2015; Stuart et al., 2019). Dead or apoptotic cells were removed if >7% of UMIs mapping to mitochondrial genes. Any cells with <200 or >9000 genes were also removed before downstream analysis. The ‘ElbowPlot’ function and principal component ‘Heatmaps’ helped determine how many principal components were used for unsupervised cluster modeling. Clustering calculations used linear dimensional reduction and visualization was done using non-linear dimensional reductions, uniform manifold projection (UMAP). 40 principal components were used for calculations. 21 principal components and 40 principal components were used for the epithelial subset and the stroma subset calculations, respectively. The genes defining the unsupervised clustering were determined using the function ‘FindAllMarkers.’ Visualization in R required library packages cowplot v1, dplyr v0.8.5, ggplot2 v3.3.0, patchwork v1, magrittr v1.5, and stringr v1.4. Stromal subpopulation analysis in Metascape (Zhou et al., 2019) used all positively differentiated genes in *Pdgfrα^+^* stromal clusters.

### 7.4 E16 and adult gland cryopreservation and sectioning

E16 and adult salivary glands were removed from CD-1 mice and preserved following a standard method (Shubin et al., 2017). Gland were fixed with 4% paraformaldehyde (Electron Microscopy Sciences #15710) in 1xPBS for 2 hours. The fixative was washed 3 times using 1xPBS. The glands were dehydrated using 3 consecutive 1-hour sucrose (Fisher #57-50-1) incubations at 4°C, at 5%, 10%, and 15%. Followed by 2 overnight incubations at 4°C first with 30%, then 15% sucrose and 50% Tissue Freezing Medium (TFM) (Electron Microscopy Sciences #72592). The fixed and dehydrated glands were frozen in TFM indirectly over liquid nitrogen. Cryosections were cut at 10 µM using Leica CM1860 and adhered to charged histological slides (Electron Microscopy Sciences #71869-10).

### 7.5 E16 PDGFRα^+^ Stromal cell isolation

Enriched E16 SMG stroma from the SMG isolation were pelleted at 300xG for 8 minutes. The media was removed, and the stroma was resuspended in 90 µL FACS buffer ((1xHank’s buffered saline solution (HBSS) (Life Technologies #14175095), 10% FBS, 5 mM Ethylenediaminetetraacetic acid (EDTA) (Falcon #S311-100), 5 mM ethylene glycol-bis(β-aminoethyl ether)-N,N,N′,N′-tetra acetic acid (EGTA) (Sigma #E4378-10G)) and positively selected using 10 uL PDGFRα antibody-coated microbeads (1:10; Miltenyi Biotec #130-101-547). PDGFRα^+^ stoma was placed into magnetic separation MS columns (Miltenyi Biotec #130-042-021) and isolated following the manufacturer’s instructions. PDGFRα^+^ stoma was pelleted 300xG for 8 minutes, resuspended in 1 mL of media, and cells counted using a hemocytometer.

### 7.6 Organoid culture

∼900 Epithelial clusters (derived from 1 gland) grown with or without 20,000-50,000 stromal cells (derived from 0.2 glands) were embedded in Matrigel (Corning #356234) at a 1:1 ratio and seeded on 0.1 μm pore porous polycarbonate filters (Nuclepore Whatman #0930051) floated on-top of media in 50 mm glass-bottom dishes (MatTek P50G-1.5-14F). The cell preparations were incubated at 37°C in a tissue culture incubator (ThermoFisher Scientific Forma Series II) with 5% CO2 for 7 days to form organoids. Media was replaced after incubating for 4 days. Media used in these experiments included DMEM/F12 with 10% FBS and Pen-Strep with growth factors or inhibitors added. Growth factors included: Amphiregulin (AREG, 100 ng/mL; Peprotech #315-36), Bone morphogenetic protein 2 (BMP2, 100-200 ng/mL; Peprotech #120-02), Bone morphogenetic protein 4 (BMP4, 100-200 ng/mL; Peprotech #315-27), Bone morphogenetic protein 7 (BMP7, 100-200 ng/mL; Peprotech #120-03P), Fibroblast growth factor-2 (FGF2, 100 ng/mL; Peprotech #450-33), Epiregulin (EREG, 100 ng/mL; Peprotech #100-04), and Transforming growth factor beta-1 (TGFβ, 5 ng/mL; Peprotech #100-12). Growth factors were solubilized in 0.2% Bovine Serum Albumin (BSA) (Sigma Millipore #A2934-100G) and stored at -20°C in single-use aliquots prior to use in epithelial cell cultures. Chemical inhibitor CHIR99021 (10 µM; Sigma Millipore #SML1046) was concentrated to 1000x in DMSO and stored at -20°C.

### 7.7 2D Stromal cell culture

Primary E16 SMG stromal cells were cultured in 10 cm tissue culture dishes (Corning #430167) containing 10 mL DMEM/F12 + 10% FBS + PenStrep with media replenished after plating and then every two days. Cells were incubated at 37°C with 5% CO2 and passed when the density reached 90%. Cells were passaged and isolated using standard trypsinization (Gibco #25200056) technique. For growth factor treatments, 50,000 SMG stromal cell (about 0.2 glands worth of cells) at passage 1 were seeded on-top of 50 mm glass bottom dishes (MatTek P50G-1.5-14F) on day 0 and growth factors or pharmacological inhibitors added from days 1-4.

### 7.8 Organoid epithelial and stromal cell isolation for microarray

In organoid cultures, the media under the filter was replaced with 4°C FACS buffer. The FACS buffer was replaced with FACS buffer on top of the organoids, which were incubated at 4°C for 20 minutes, triturated again and collected into a 15 mL conical tube containing an equal volume of 4°C FACS buffer. The cells were triturated at 4°C to separate the cells and dissolve the Matrigel. Epithelial clusters were collected by centrifuging at 10xG for 1 minute and resuspended in 4°C FACS buffer. The single stromal cells were pelleted at 450xG for 5 minutes then resuspended in 4°C FACS buffer. The stroma was negatively selected against epithelium using a EPCAM antibody coated microbeads (1:10; Miltenyi Biotec #130-061-101) and a Miltenyi MS column.

### 7.9 RNA isolation for microarray and transcriptomic analysis

Total RNA was isolated using RNeasy mini kits (Qiagen #74134) or RNAqueous micro kits (ThermoFisher Scientific #AM1931). RNA quality was analyzed using UV/Vis Spectrophotometer (NanoDrop ND-1000) with 270/280 between 1.7-2.2 and bioanalyzer with a RIN score >7.0 as cut offs. The RNA for transcriptome analyses were prepared with Clariom S (Epithelium) (ThermoFisher Scientific #902930) or Clariom D (Stroma) (ThermoFisher Scientific #902513) mouse microarrays. Data was analyzed using Transcriptome Analysis Console v.4.0.2.15, R functions and the Broads Institute’s Gene Set Enrichment Analysis (GSEA) software v4.

### 7.10 Immunocytochemistry and imaging

SMG organoid and cell cultures grown on Nuclepore filters (Cytiva #110405) or glass were fixed with 4% paraformaldehyde (Electron Microscopy Sciences #15710) in 1xPBS for 20 minutes or in -20°C methanol for 18 minutes. Downstream antibody staining determined fixation method. Immunocytochemistry (ICC) was performed as described previously (Daley et al., 2009; Sequeira et al., 2012), with the use of 0.4% Triton-X 100 (Sigma #T9284-100ML) instead of 0.1% Triton-X for PFA-fixed samples. All primary antibodies incubations were overnight at 4°C. Secondary antibody incubations were 1-3 hours at room temperature. Primary antibodies and dilutions used include: Aquaporin 5 (AQP5) (1:400; Alomone #AQP-005), Calponin 1 (CNN1) (1:600; Abcam # ab46794), Cytokeratin 7 (KRT7) (1:200; Abcam #ab9021), EPCAM-647 (1:400; Biolegend #118212), PDGFRα (1:200; ThermoFisher Scientific #14-1401-81), PDGFRα (1:100; R&D Systems #AF1062-SP), PDGFRβ (1:200; Abcam #ab32570), PDGFRβ (1:100; ThermoFisher Scientific #ab32570), Perlecan (HSPG2) (1:200; SantaCruz #sc-33707), Smooth Muscle Actin Alpha (αSMA/ACTA2) (1:1000; Sigma #12-1402-81), THY1 (1:200; Biolegend #105301), Endoglin (ENG) (1:100; ThermoFisher Scientific # 14-1051-81), and Vimentin (VIM) (1:2000; Sigma #V2258). Secondary antibodies included Cyanine and Alexa dye-conjugated AffiniPure F(ab’^’^)2 fragments purchased from Jackson ImmunoResearch Laboratories and used at a dilution of 1:250. 4′,6-diamidino-2-phenylindole (DAPI) (Life Technologies #D1306) was used for nuclei staining in conjunction with secondary antibodies. Nuclepore filters were mounted using antifade mounting media containing 90% (vol/vol) glycerol (Sigma #G5516-1L) in 1xPBS with 1-4 diazobicyclo[2,2,2]octane (Sigma #D27802-100G) and n-propyl gallate (Sigma #P3130-100G) as antifade agents (Gerdes et al., 2013; Valnes & Brandtzaeg, 1985). Imaging was performed using Zeiss Z1 Cell Observer widefield, Zeiss LSM 710 confocal, or the Leica TCS SP5 confocal microscope at 10x, 20x, and 63x (oil immersion) magnifications with the same microscope and laser configurations for all samples within an experiment, as appropriate to the imaging method.

### 7.11 Quantification and Statistical analyses

SEURAT v3.1.5 (Satija et al., 2015; Stuart et al., 2019) uses a non-parametric Wilcoxon rank sum tests for statistical significance. Microarrays used RMA normalization and log 2 transform in Transcriptome Analysis Console v4.0.2 (ThermoFisher Scientific) or in R (R Core Team, 2021). Adjusted p-values of <0.05 and log fold change >1.99 were parameters used to determine differentially expressed genes. For immunocytochemistry images, FIJI v1.53c performed the rolling-ball background subtraction then color thresholding was used to calculate staining area. Staining area for organoid stromal measurements used ROIs drawn by hand to exclude the epithelial components. Staining area from organoid epithelial measurements used color thresholding on DAPI to exclude any stromal staining. At least one representative image from each sample was taken at 5x widefield magnification that framed 20-40% of the entire sample. All statistical significances calculated between stained areas were done using a Student’s T-test for dual-sample comparison or single-factor ANOVA followed with post-hoc Tukey HSD test for multi-sample comparison. Statistical tests were performed in Microsoft Excel (Microsoft Corporation) or in R. Staining area results are expressed as mean ± standard deviation (s.d.).

### 7.12 Sequencing and Microarray Data Availability

Raw data from both our scRNA Seq and microarray studies can be found on the Gene Expression Omnibus (GEO) repository. All datasets are listed under SuperSeries GSE181430 or individually as GSE181425 (scRNA Seq) and GSE181423, GSE181424 (Clariom S and D microarrays). All these data will be released by GEO on 01/30/2022. Scripts used to analyze data can be found on github: https://github.com/MLarsenLab

### 8.0 Acknowledgements

The authors would like to thank University at Albany’s Core facilities and the Center for Functional Genomics for equipment and expertise.

---

**Supplemental Figure 1:**
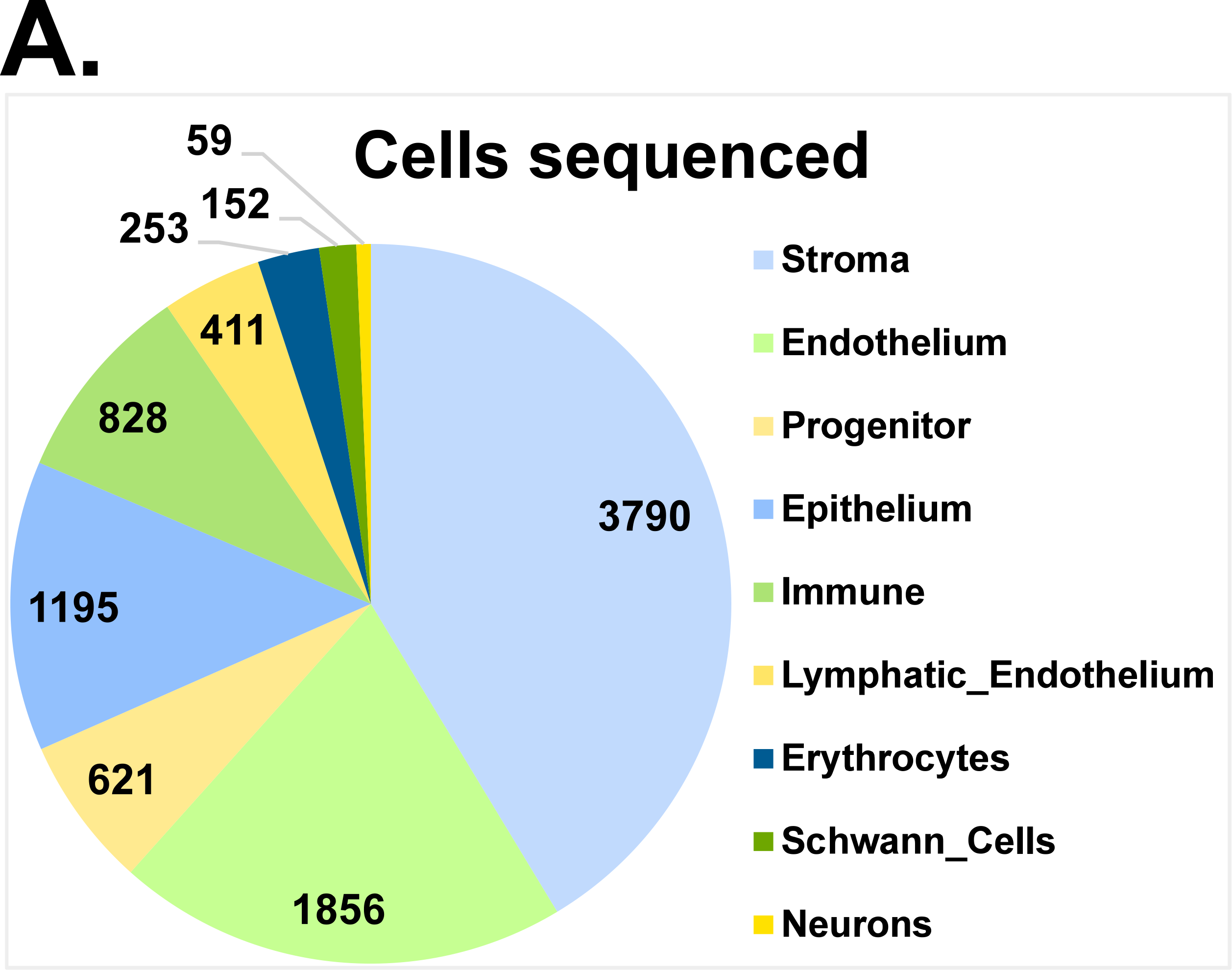
scRNA sequencing 1125 cell composition. A) Pie chart showing proportions of cells sequenced based on Seurat Clustering.

**Supplemental Figure 2:**
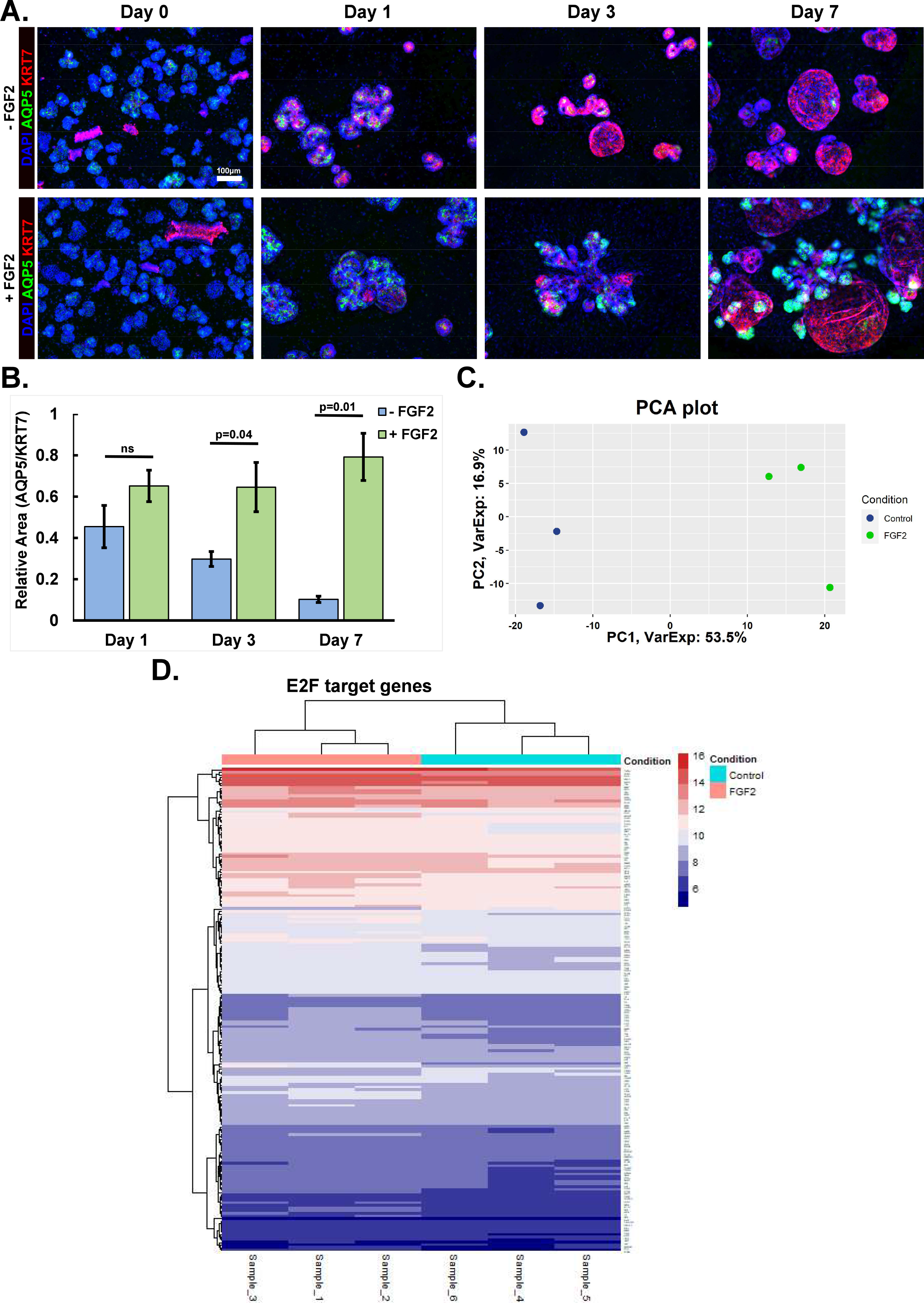
FGF2 signaling in stroma induced acinar phenotype in salivary gland organoids. A) E16 epithelial and stromal cell-containing organoids were grown for 0, 1, 3, and 7 days. Organoid epithelium grown without FGF2 have ductal phenotype and are Cytokeratin 7^+^ (KRT7, red), while FGF2 cultured organoid epithelium have a proacinar phenotype and are Aquaporin 5^+^ (AQP5, green). Based on immunocytochemistry (ICC) with DAPI (blue) nuclear stain. Scale: 100 µm. B) ICC area quantification showing the AQP5/KRT7 ratio increasing over time; n=4 technical replicates, Two-tailed Unpaired Student T-Test with p-value indicated. C) PCA plot comparing Clariom S microarray data generated from epithelial organoids grown with or without FGF2. More than half of the variance is described by the first principal component; n=3 experimental replicates. D) Transcriptome heatmap showing E2F gene enrichment of FGF2 (lavender) and control (cyan) organoid culture conditions. Log two-fold scale for gene expression is shown with red (higher) and blue (lower) shown.

**Supplemental Figure 3:**
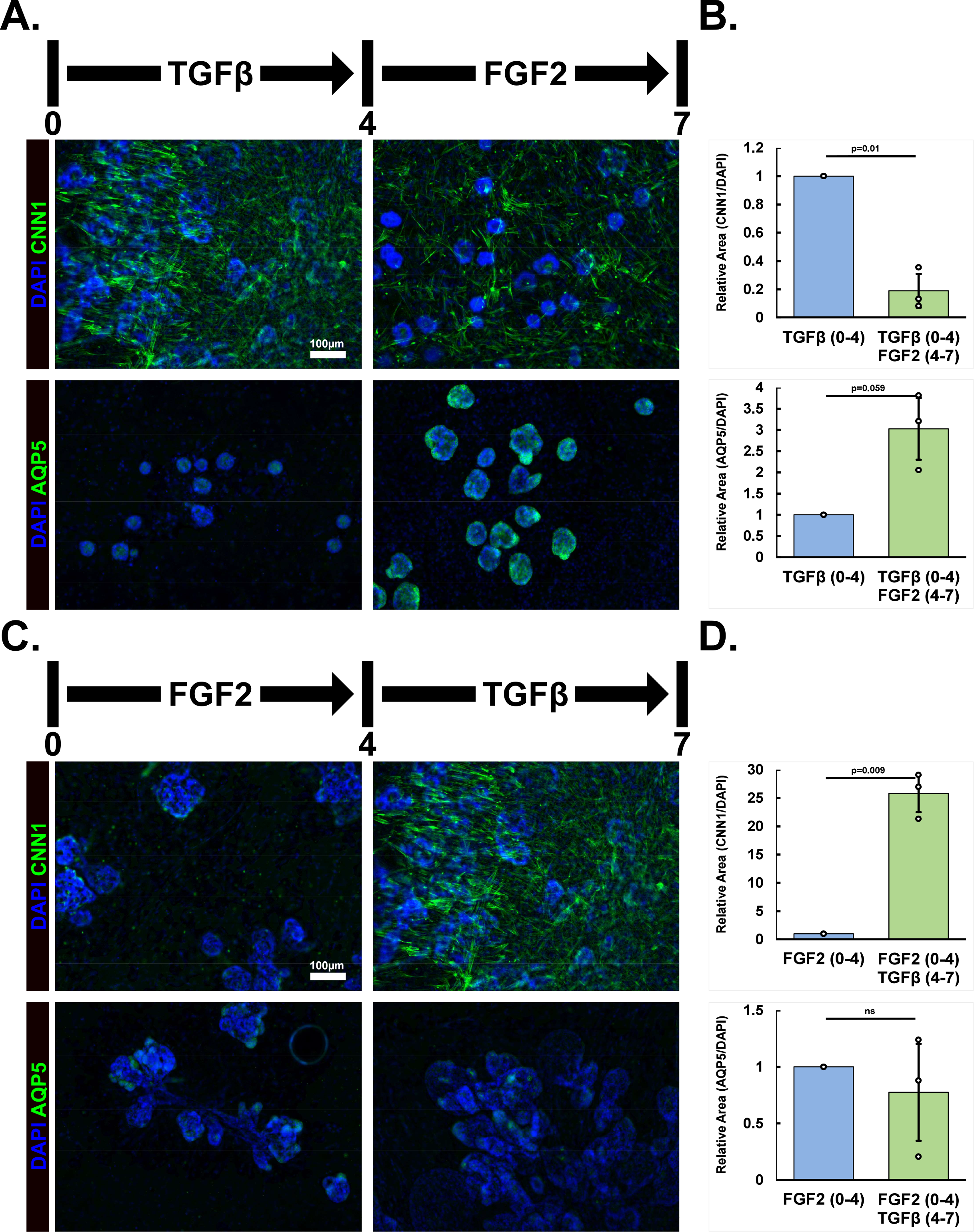
TGFβ signaling correlates myofibroblast differentiation and proacinar dedifferentiation. A) Organoids growth with TGFβ1 containing media for 4 followed by FGF2 containg media for 3 days. The FGF2 media change reduced myofibroblast marker staining but increases proacinar marker staining. Scale: 100 µm. B) Relative staining of AQP5 or CNN1 normalized to DAPI. Myofibroblast ICC staining decreases 5-fold, while proacinar staining increases 3-fold; n=3 technical replicates, Two-tailed Unpaired Student t-test. C) Organoids growth with FGF2 containing media 4 days followed by TGFβ1 containing  media for 3 days. The TGFβ1 media change increased myofibroblast marker staining and decreased proacinar marker staining. Scale: 100 µm. D) Relative staining AQP5 or CNN1 normalized to DAPI. Myofibroblast ICC staining increases 25-fold, while proacinar staining remained unchanged; n=3 technical replicates, Two-tailed Unpaired Student t-test.

